# Non-essential function of KRAB zinc finger gene clusters in retrotransposon suppression

**DOI:** 10.1101/2020.01.17.910679

**Authors:** Gernot Wolf, Alberto de Iaco, Ming-An Sun, Melania Bruno, Matthew Tinkham, Don Hoang, Apratim Mitra, Sherry Ralls, Didier Trono, Todd S. Macfarlan

## Abstract

The Krüppel-associated box zinc finger protein (KRAB-ZFP) family amplified and diversified in mammals by segmental duplications, but the function of the majority of this gene family remains largely unexplored due to the inaccessibility of the gene clusters to conventional gene targeting. We determined the genomic binding sites of 61 murine KRAB-ZFPs and genetically deleted in mouse embryonic stem (ES) cells five large KRAB-ZFP gene clusters encoding nearly one tenth of the more than 700 mouse KRAB-ZFPs. We demonstrate that clustered KRAB-ZFPs directly bind and silence retrotransposons and block retrotransposon-borne enhancers from gene activation in ES cells. Homozygous knockout mice generated from ES cells deleted in one of two KRAB-ZFP clusters were born at sub-mendelian frequencies in some matings, but heterozygous intercrosses could also yield knockout progeny with no overt phenotype. We further developed a retrotransposon capture-sequencing approach to assess mobility of the MMETn family of endogenous retrovirus like elements, which are transcriptionally activated in KRAB-ZFP cluster KOs, in a pedigree of KRAB-ZFP cluster KO and WT mice. We identified numerous somatic and several germ-line MMETn insertions, and found a modest increase in activity in mutant animals, but these events were detected in both wild-type and KO mice in stochastic and highly variable patterns. Our data suggests that the majority of young KRAB-ZFPs play a non-essential role in transposon silencing, likely due to the large redundancy with other KRAB-ZFPs and other transposon restriction pathways in mice.

**One Sentence Summary:** Megabase-scale deletions of KRAB-ZFP gene clusters in mice leads to retrotransposon activation.

## Introduction

Nearly half of the human and mouse genomes consist of transposable elements (TEs). TEs shape the evolution of species, serving as a source for genetic innovation (*1, 2*). However, TEs also potentially harm their hosts by insertional mutagenesis, gene deregulation and activation of innate immunity (*3-6*). To protect themselves from TE activity, host organisms have developed a wide range of defense mechanisms targeting virtually all steps of the TE life cycle (*7*). In tetrapods, KRAB zinc finger protein (KRAB-ZFP) genes have amplified and diversified, likely in response to TE colonization (*8-11*). Conventional ZFPs bind DNA using tandem arrays of C2H2 zinc finger domains, each capable of specifically interacting with three nucleotides, whereas some zinc fingers can bind two or four nucleotides and include DNA backbone interactions depending on target DNA structure (*12*). This allows KRAB-ZFPs to flexibly bind to large stretches of DNA with high affinity. The KRAB domain binds the corepressor KAP1, which in turn recruits histone modifying enzymes including the NuRD histone deacetylase complex and the H3K9-specific methylase SETDB1 (*13, 14*), which induces persistent and heritable gene silencing (*15*). Deletion of KAP1 (*16*) or SETDB1 (*17*) in mouse embryonic stem (ES) cells induces TE reactivation and cell death, but only minor phenotypes in differentiated cells, suggesting KRAB-ZFPs are most important during early embryogenesis where they mark TEs for stable epigenetic silencing that persists through development. However, SETDB1-containing complexes are also required to repress TEs in primordial germs cells (*18*) and adult tissues (*19*), indicating KRAB-ZFPs are active beyond early development.

TEs, especially long terminal repeat (LTR) retrotransposons, also known as endogenous retroviruses (ERVs), can affect expression of neighboring genes through their promoter and enhancer functions (*20-22*). KAP1 deletion in mouse ES cells causes rapid gene deregulation (*23*), indicating that KRAB-ZFPs may regulate gene expression by recruiting KAP1 to TEs. Indeed, *Zfp809* knock-out (KO) in mice resulted in transcriptional activation of a handful of genes in various tissues adjacent to ZFP809-targeted VL30-Pro elements (*24*). It has therefore been speculated that KRAB-ZFPs bind to TE sequences to domesticate them for gene regulatory innovation (*25*). This idea is supported by the observation that many human KRAB-ZFPs target TE groups that have lost their coding potential millions of years ago and that KRAB-ZFP target sequences within TEs are in some cases under purifying selection (*8*). However, there are also clear signs of an evolutionary arms-race between human TEs and KRAB-ZFPs (*26*), indicating that some KRAB-ZFPs may limit TE mobility for stretches of evolutionary time, prior to their ultimate loss from the genome or adaptation for other regulatory functions. Here we use the laboratory mouse, which has undergone a recent expansion of the KRAB-ZFP family, to determine the *in vivo* requirement of the majority of evolutionarily young KRAB-ZFP genes.

## Results

### Mouse KRAB-ZFPs target retrotransposons

We analyzed the RNA expression profiles of mouse KRAB-ZFPs across a wide range of tissues to identify candidates active in early embryos/ES cells. While the majority of KRAB-ZFPs are expressed at low levels and uniformly across tissues, a group of KRAB-ZFPs are highly and almost exclusively expressed in ES cells (Supplemental Fig. 1A). About two thirds of these KRAB-ZFPs are physically linked in two clusters on chromosome 2 (Chr2-cl) and 4 (Chr4-cl) (Supplemental Fig. 1B). These two clusters encode 40 and 21 KRAB-ZFP annotated genes, respectively, which, with one exception on Chr4-cl, do not have orthologues in rat or any other sequenced mammals (Supplemental Table 1). The KRAB-ZFPs within these two genomic clusters also group together phylogenetically (Supplemental Fig. 1C), indicating these gene clusters arose by a series of recent segmental gene duplications (*27*).

**Figure 1.**
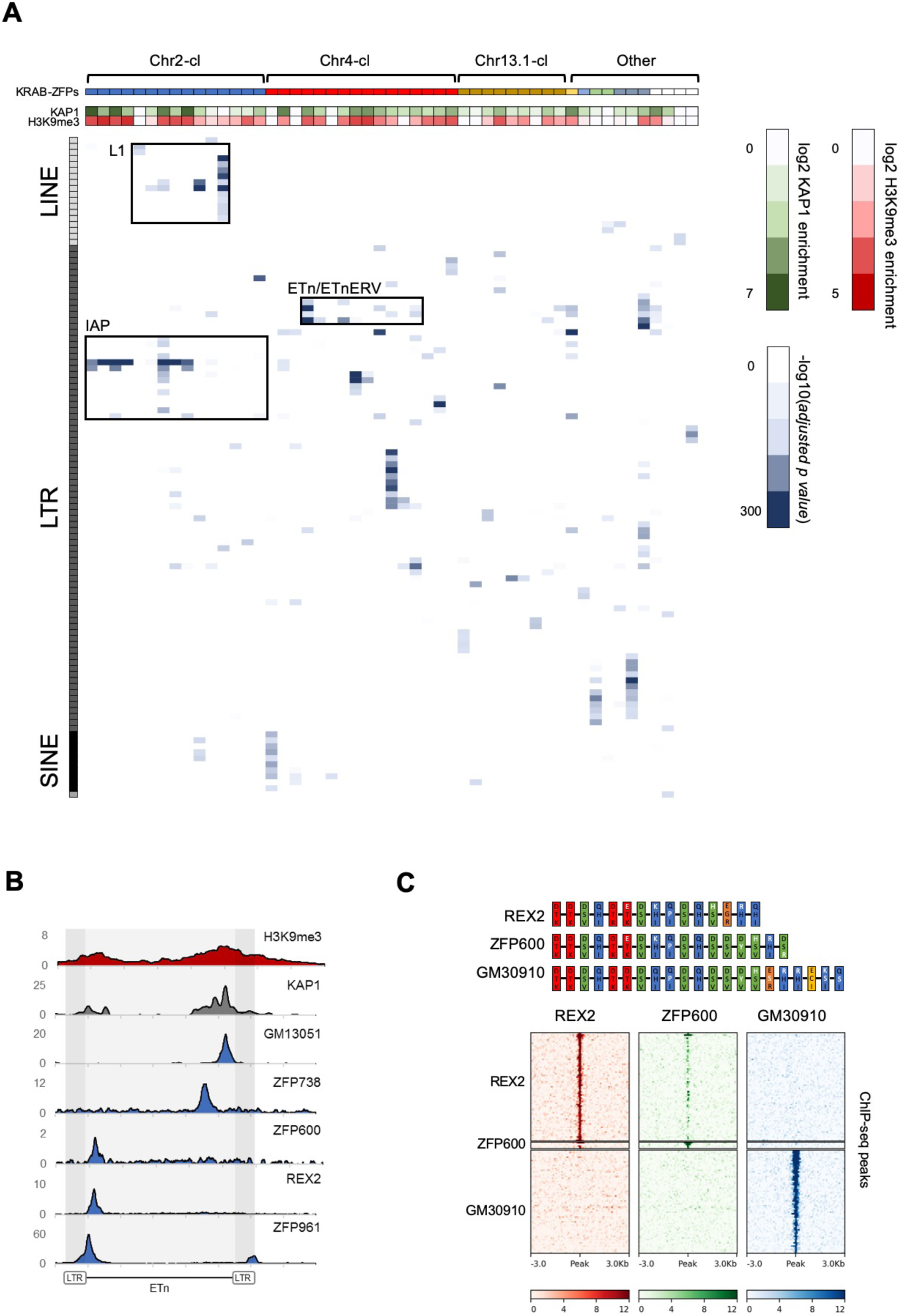
Genome-wide binding patterns of mouse KRAB-ZFPs. (A) Probability heatmap of KRAB-ZFP binding to TEs. Blue color intensity (main field) corresponds to -log10 [*adjusted p-value*] enrichment of ChIP-seq peak overlap with TE groups (Fisher’s exact test). The green/red color intensity (top panel) represent mean KAP1 (GEO accession: GSM1406445) and H3K9me3 (GEO accession: GSM1327148) enrichment (respectively) at peaks overlapping significantly targeted TEs (*adjusted p-value* < 1e-5) in WT ES cells. (B) Summarized ChIP-seq signal for indicated KRAB-ZFPs and previously published KAP1 and H3K9me3 in WT ES cells across 127 intact ETn elements. (C) Heatmaps of KRAB-ZFP ChIP-seq signal at ChIP-seq peaks. For better comparison, peaks for all three KRAB-ZFP were called with the same parameters (*P* < 1e-10, peak enrichment > 20). The top panel shows a schematic of the arrangement of the contact amino acid composition of each zinc finger. Zinc fingers are grouped and colored according to similarity, with amino acid differences relative to the five consensus fingers highlighted in white.

To determine the binding sites of the KRAB-ZFPs within these and other gene clusters, we expressed epitope-tagged KRAB-ZFPs using stably integrating vectors in mouse embryonic carcinoma (EC) or ES cells (Table 1, Supplemental Table 1) and performed chromatin immunoprecipitation followed by deep sequencing (ChIP-seq). We then determined whether the identified binding sites are significantly enriched over annotated TEs and used the non-repetitive peak fraction to identify binding motifs. We discarded 7 of 68 ChIP-seq datasets because we could not obtain a binding motif or a target TE and manual inspection confirmed low signal to noise ratio. Of the remaining 61 KRAB-ZFPs, 51 significantly overlapped at least one TE subfamily (*adjusted p value* < 1e-5). Altogether, 81 LTR retrotransposon, 18 LINE, 10 SINE and one DNA transposon subfamilies were targeted by at least one of the 51 KRAB-ZFPs (Supplemental Table 1 and Supplemental Table 2). Chr2-cl KRAB-ZFPs preferably bound IAPEz retrotransposons and L1-type LINEs, while Chr4-cl KRAB-ZFPs targeted various retrotransposons, including MMETn (hereafter referred to as ETn) and their coding counterparts, ETnERV (Fig. 1A and Supplemental Table 2). The validity of our ChIP-seq screen was confirmed by the identification of binding motifs - which often resembled the computationally predicted motifs (Supplemental Fig. 2A) - for the majority of screened KRAB-ZFPs (Supplemental Table 1). Moreover, predicted and experimentally determined motifs were found in targeted TEs in most cases (Supplemental Table 1), and reporter repression assays confirmed KRAB-ZFP induced silencing for all the tested sequences (Supplemental Fig. 2B). Finally, we observed KAP1 and H3K9me3 enrichment at most of the targeted TEs in wild type ES cells, indicating that most of these KRAB-ZFPs are functionally active in the early embryo (Fig. 1A).

**Table 1:** KRAB-ZFP genes clusters in the mouse genome that were investigated in this study. * Number of protein-coding KRAB-ZFP genes identified in a previously published screen (*8*) and the ChIP-seq data column indicates the number of KRAB-ZFPs for which ChIP-seq was performed in the this study.

| Cluster | Location | Size (Mb) | # of KRAB-ZFPs* | ChIP-seq data |
| --- | --- | --- | --- | --- |
| Chr2 | Chr2 qH4 | 3.1 | 40 | 17 |
| Chr4 | Chr4 qE1 | 2.3 | 21 | 19 |
| Chr10 | Chr10 qC1 | 0.6 | 6 | 1 |
| Chr13.1 | Chr13 qB3 | 1.2 | 6 | 2 |
| Chr13.2 | Chr13 qB3 | 0.8 | 26 | 12 |
| Chr8 | Chr8 qB3.3 | 0.1 | 4 | 4 |
| Chr9 | Chr9 qA3 | 0.1 | 4 | 2 |
| Other | - | - | 248 | 4 |

**Figure 2.**
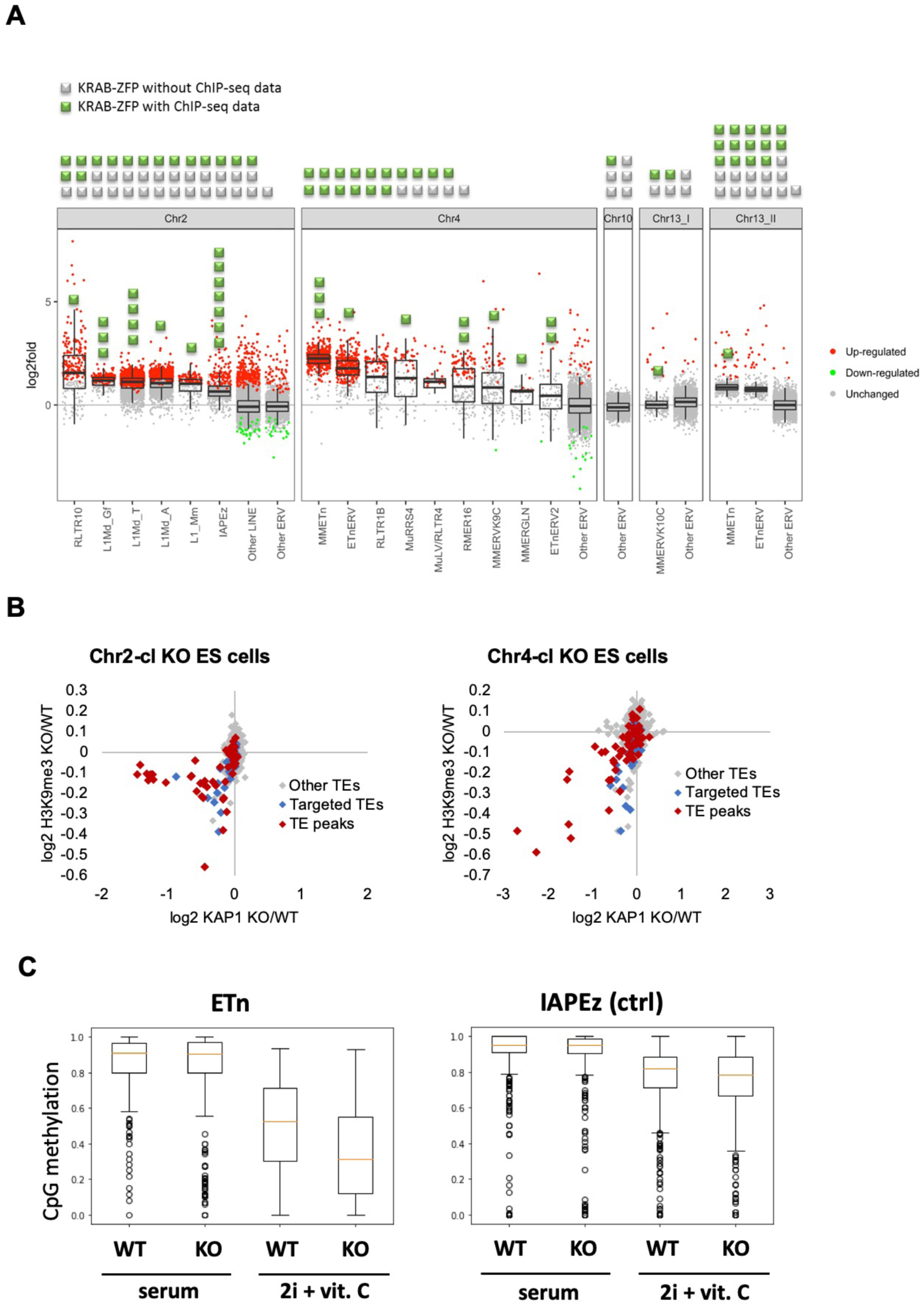
Retrotransposon reactivation in KRAB-ZFP cluster KO ES cells. (A) RNA-seq analysis of TE expression in five KRAB-ZFP cluster KO ES cells. Green and grey squares on top of the panel represent KRAB-ZFPs with or without ChIP-seq data, respectively, within each deleted gene cluster. Reactivated TEs that are bound by one or several KRAB-ZFPs are indicated by green squares in the panel. Significantly up- and downregulated elements (*adjusted p-value* < 0.05) are highlighted in red and green, respectively. (B) Differential KAP1 binding and H3K9me3 enrichment at TE groups (summarized across all insertions) in Chr2-cl and Chr4-cl KO ES cells. TE groups targeted by one or several KRAB-ZFPs encoded within the deleted clusters are highlighted in blue (differential enrichment over the entire TE sequences) and red (differential enrichment at TE regions that overlap with KRAB-ZFP ChIP-seq peaks). (C) DNA methylation status of CpG sites at indicated TE groups in WT and Chr4-cl KO ES cells grown in serum containing media or in hypomethylation-inducing media (2i + Vitamin C).

We generally observed that KRAB-ZFPs present exclusively in mouse target TEs that are restricted to the mouse genome, indicating KRAB-ZFPs and their targets emerged together. For example, several mouse-specific KRAB-ZFPs in Chr2-cl and Chr4-cl target IAP and ETn elements which are only found in the mouse genome and are highly active. This is the strongest data to date supporting that recent KRAB-ZFP expansions in these young clusters is a response to recent TE activity. Likewise, ZFP599 and ZFP617, both conserved in Muroidea, bind to various ORR1-type LTRs which are present in the rat genome (Supplemental Table 1). However, ZFP961, a KRAB-ZFP encoded on a small gene cluster on chromosome 8 that is conserved in Muroidea targets TEs that are only found in the mouse genome (e.g. ETn), a paradox we have previously observed with ZFP809, which also targets TEs that are evolutionarily younger than itself (*24*). The ZFP961 binding site is located at the 5’ end of the internal region of ETn and ETnERV elements, a sequence that usually contains the primer binding site (PBS), which is required to prime retroviral reverse transcription. Indeed, the ZFP961 motif closely resembles the PBS^Lys1,2^ (Supplemental Fig. 3A), which had been previously identified as a KAP1-dependent target of retroviral repression (*28, 29*). Repression of the PBS^Lys1,2^ by ZFP961 was also confirmed in reporter assays (Supplemental Fig. 2B), indicating that ZFP961 is likely responsible for this silencing effect.

**Figure 3.**
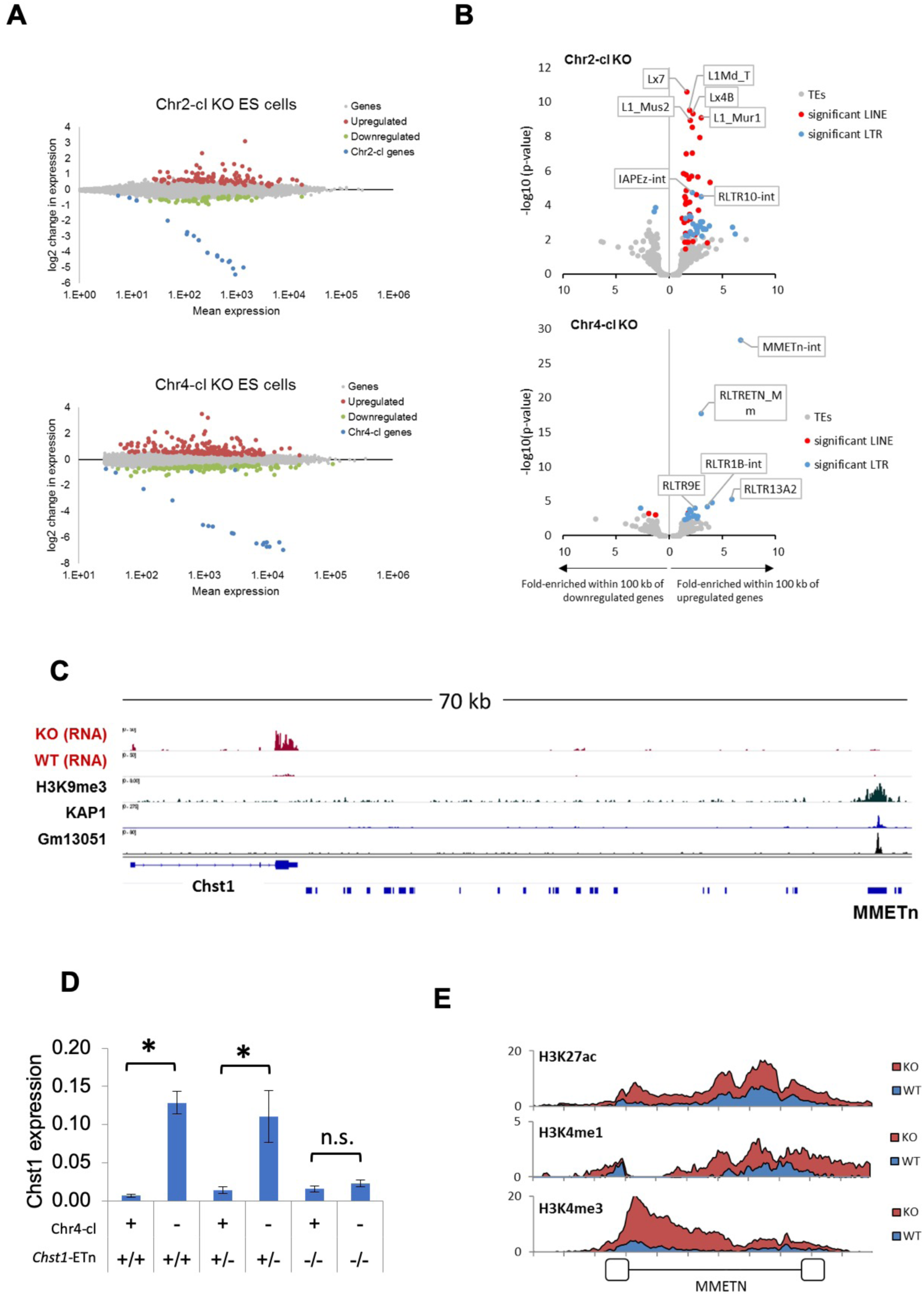
TE-dependent gene activation in KRAB-ZFP cluster KO ES cells. (A) Differential gene expression in Chr2-cl and Chr4-cl KO ES cells. Significantly up- and downregulated genes (*adjusted p-value* < 0.05) are highlighted in red and green, respectively, KRAB-ZFP genes within the deleted clusters are shown in blue. (B) Correlation of TEs and gene deregulation. Plots show enrichment of TE groups within 100 kb of up- and downregulated genes relative to all genes. Significantly overrepresented LTR and LINE groups (*adjusted p-value* < 0.1) are highlighted in blue and red, respectively. (C) Schematic view of the downstream region of *Chst1* where a 5’truncated ETn insertion is located. ChIP-seq (Input subtracted from ChIP) data for overexpressed epitope-tagged Gm13051 (a Chr4-cl KRAB-ZFP) in F9 EC cells, and re-mapped KAP1 (GEO accession: GSM1406445) and H3K9me3 (GEO accession: GSM1327148) in WT ES cells are shown together with RNA-seq data from Chr4-cl WT and KO ES cells (mapped using Bowtie (-a -m 1 -- strata -v 2) to exclude reads that cannot be uniquely mapped). (D) RT-qPCR analysis of Chst1 mRNA expression in Chr4-cl WT and KO ES cells with or without the CRISPR/Cas9 deleted ETn insertion near Chst1. Values represent mean expression (normalized to Gapdh) from three biological replicates per sample (each performed in three technical replicates) in arbitrary units. Error bars represent standard deviation and asterisks indicate significance (*p* < 0.01, Student’s t-test). n.s.: not significant. (E) Mean coverage of ChIP-seq data (Input subtracted from ChIP) in Chr4-cl WT and KO ES cells over 127 full-length ETn insertions.

Using previously generated ETn and ETnERV retrotransposition reporters in which we mutated KRAB-ZFP binding sties (*30*), we tested and confirmed that the REX2/ZFP600 and GM13051 binding sites within these TEs are required for efficient retrotransposition (Supplemental Fig. 3B). REX2 and ZFP600 both bind a target about 200 bp from the start of the internal region (Fig. 1B), a region that often encodes the packaging signal. GM13051 binds a target coding for part of a highly structured mRNA export signal (*31*) near the 3’end of the internal region of ETn (Supplemental Fig. 3C). Both signals are characterized by stem-loop intramolecular base-pairing in which a single mutation can disrupt loop formation. This indicates that at least some KRAB-ZFPs evolved to bind functionally essential target sequences which cannot easily evade repression by mutation.

Our KRAB-ZFP ChIP-seq dataset also provided unique insights into the emergence of new KRAB-ZFPs and binding patterns. The Chr4-cl KRAB-ZFPs REX2 and ZFP600 bind to the same target within ETn but with varying affinity (Fig. 1C). Comparison of the amino acids responsible for DNA contact revealed a high similarity between REX2 and ZFP600, with the main differences at the most C-terminal zinc fingers. Additionally, we found that GM30910, another KRAB-ZFP encoded in the Chr4-cl, also shows a strong similarity to both KRAB-ZFPs yet targets entirely different groups of TEs (Fig. 1C and Supplemental Table 2). Together with previously shown data (*19*), this example highlights how addition of a few new zinc fingers to an existing array can entirely shift the mode of DNA binding.

### Genetic deletion of KRAB-ZFP gene clusters leads to retrotransposon reactivation

The majority of KRAB-ZFP genes are harbored in large, highly repetitive clusters that have formed by successive complex segmental duplications (*27*), rendering them inaccessible to conventional gene targeting. We therefore developed a strategy to delete entire KRAB-ZFP gene clusters in ES cells (including the Chr2-cl and Chr4-cl as well as two clusters on chromosome 13 and a cluster on chromosome 10) using two CRISPR/Cas9 gRNAs targeting unique regions flanking each cluster, and short single-stranded repair oligos with homologies to both sides of the projected cut sites. Using this approach, we generated five cluster KO ES cell lines in at least two biological replicates and performed RNA sequencing (RNA-seq) to determine TE expression levels. Strikingly, four of the five cluster KO ES cells exhibited distinct TE reactivation phenotypes (Fig. 2A). Chr2-cl KO resulted in reactivation of several L1 subfamilies as well as RLTR10 and IAPEz ERVs. In contrast, the most strongly upregulated TEs in Chr4-cl KO cells were ETn/ETnERV, with several other ERV groups modestly reactivated. ETn/ETnERV elements were also upregulated in Chr13.2-cl KO ES cells while the only upregulated ERV in Chr13.1-cl KO ES cells were MMERVK10C elements (Fig. 2A). Most reactivated retrotransposons were targeted by at least one KRAB-ZFP that was encoded in the deleted cluster (Fig. 2A and Supplemental Table 1), indicating a direct effect of these KRAB-ZFPs on TE expression levels. Furthermore, we observed a loss of KAP1 binding and H3K9me3 at several TE subfamilies that are targeted by at least one KRAB-ZFP within the deleted Chr2-cl and Chr4-cl (Fig. 2B, Supplemental Fig. 4A and Supplemental Table 3), including L1, ETn and IAPEz elements. Using reduced representation bisulfite sequencing (RRBS-seq), we found that a subset of KRAB-ZFP bound TEs were partially hypomethylated in Chr4-cl KO ESCs, but only when grown in genome-wide hypomethylation-inducing conditions (*32*) (Fig. 2C and Supplemental Table 4). These data are consistent with the hypothesis that KRAB-ZFPs/KAP1 are not required to establish DNA methylation, but under certain conditions they protect specific TEs and imprint control regions from genome-wide demethylation (*33, 34*).

**Figure 4.**
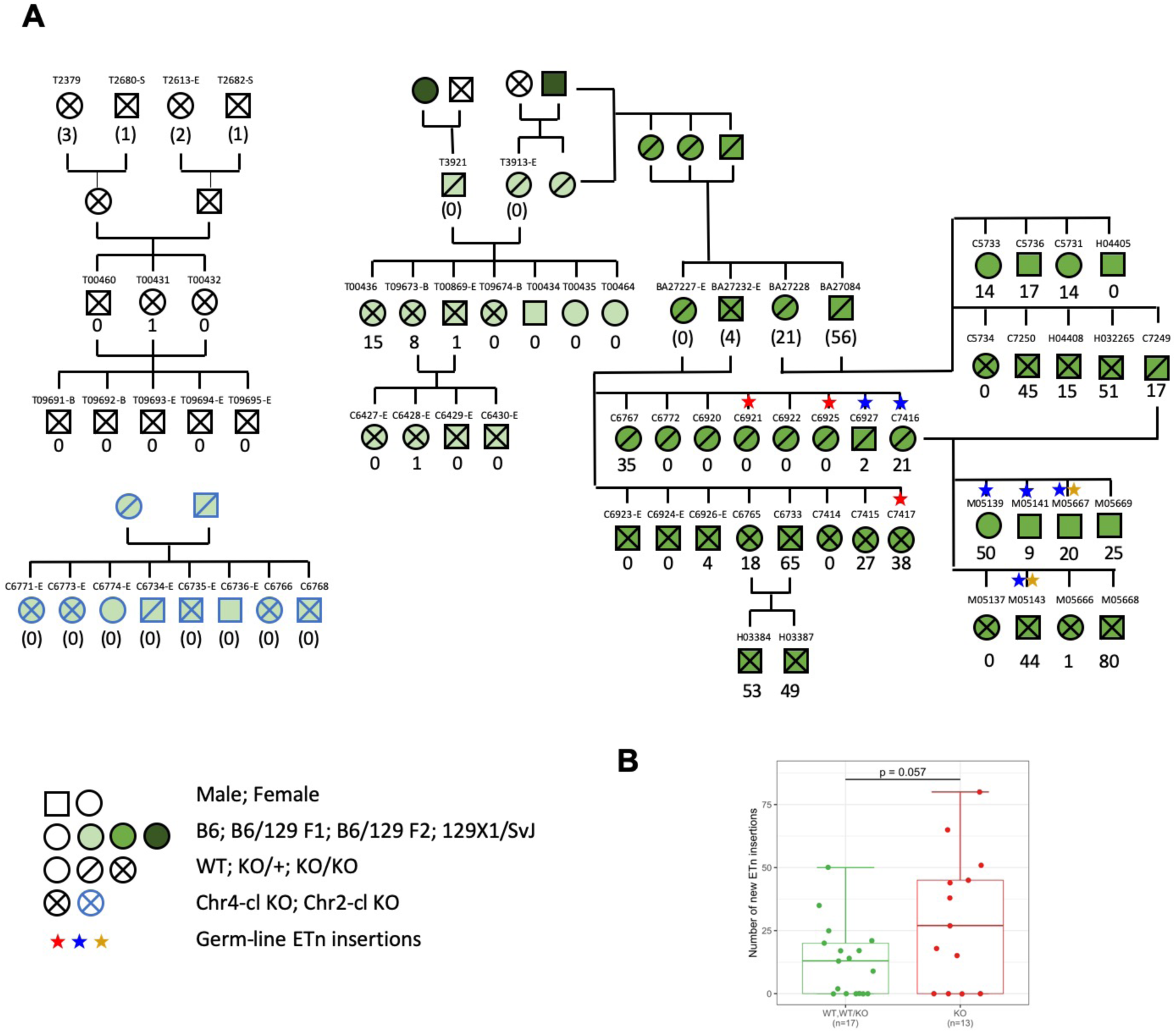
ETn retrotransposition in Chr4-cl KO mice. (A) Pedigree of mice used for transposon insertion screening by capture-seq in mice of different strain backgrounds. The number of novel ETn insertions (only present in one animal) are indicated. For animals whose direct ancestors have not been screened, the ETn insertions are shown in parentheses since parental inheritance cannot be excluded in that case. Germ-line insertions are indicated by asterisks. All DNA samples were prepared from tail tissues unless noted (-S: spleen, -E: ear, -B:Blood) (B) Statistical analysis of ETn insertion frequency in tail tissue from 30 Chr4-cl KO, KO/WT and WT mice that derived from two different Chr4-cl KO/WT x KO/WT matings. Only DNA samples that were collected from juvenile tails were considered for this analysis. P-value was calculated using one-sided Wilcoxon Rank Sum Test. WT and KO/WT mice were combined for the statistical analysis.

### KRAB-ZFP cluster deletions license TE-borne enhancers

We next used our RNA-seq datasets to determine the effect of KRAB-ZFP cluster deletions on gene expression. We identified 195 significantly upregulated and 130 downregulated genes in Chr4-cl KO ES cells, and 108 upregulated and 59 downregulated genes in Chr2-cl KO ES cells (excluding genes on the deleted cluster) (Fig. 3A). To address whether gene deregulation in Chr2-cl and Chr4-cl KO ES cells is caused by nearby TE reactivation, we determined whether genes near certain TE subfamilies are more frequently deregulated than random genes. We found a strong correlation of gene upregulation and TE proximity for several TE subfamilies, of which many became transcriptionally activated themselves (Fig. 3B). For example, nearly 10% of genes that are located within 100 kb (up-or downstream of the TSS) of an ETn element are upregulated in Chr4-cl KO ES cells, as compared to 0.8% of all genes. In Chr2-cl KO ES cells, upregulated genes were significantly enriched near various LINE groups but also IAPEz-int and RLTR10-int elements, indicating that TE-binding KRAB-ZFPs in these clusters limit the potential activating effects of TEs on nearby genes.

While we generally observed that TE-associated gene reactivation is not caused by elongated or spliced transcription starting at the retrotransposons, we did observe that the strength of the effect of ETn elements on gene expression is stronger on genes in closer proximity. About 25% of genes located within 20 kb of an ETn element, but only 5% of genes located at a distance between 50 and 100 kb from the nearest ETn insertion, become upregulated in Chr4-cl KO ES cells. Importantly however, the correlation is still significant for genes that are located at distances between 50 and 100 kb from the nearest ETn insertion, indicating that ETn elements can act as long-range enhancers of gene expression in the absence of KRAB-ZFPs that target them. To confirm that Chr4-cl KRAB-ZFPs such as GM13051 block ETn-borne enhancers, we tested the ability of a putative ETn enhancer to activate transcription in a reporter assay. For this purpose, we cloned a 5 kb fragment spanning from the GM13051 binding site within the internal region of a truncated ETn insertion to the first exon of the *Cd59a* gene, which is strongly activated in Chr4-cl KO ES cells (Supplemental Fig. 4B). We observed strong transcriptional activity of this fragment which was significantly higher in Chr4-cl KO ES cells. Surprisingly, this activity was reduced to background when the internal segment of the ETn element was not included in the fragment, suggesting the internal segment of the ETn element, but not its LTR, contains a Chr4-cl KRAB-ZFP sensitive enhancer. To further corroborate these findings, we genetically deleted an ETn element that is located about 60 kb from the TSS of *Chst1*, one of the top-upregulated genes in Chr4-cl KO ES cells (Fig. 3C). RT-qPCR analysis revealed that the *Chst1* upregulation phenotype in Chr4-cl KO ES cells diminishes when the ETn insertion is absent, providing direct evidence that a KRAB-ZFP controlled ETn-borne enhancer regulates Chst1 expression (Fig. 3D). Furthermore, ChIP-seq confirmed a general increase of H3K4me3, H3K4me1 and H3K27ac marks at ETn elements in Chr4-cl KO ES cells (Fig. 3E). Notably, enhancer marks were most pronounced around the GM13051 binding site near the 3’end of the internal region, confirming that the enhancer activity of ETn is located on the internal region and not on the LTR.

### ETn retrotransposition in Chr4-cl KO and WT mice

IAP, ETn/ETnERV and MuLV/RLTR4 retrotransposons are highly polymorphic in inbred mouse strains (*35*), indicating that these elements are able to mobilize in the germ line. Since these retrotransposons are upregulated in Chr2-cl and Chr4-cl KO ES cells, we speculated that these KRAB-ZFP clusters evolved to minimize the risks of insertional mutagenesis by retrotransposition. To test this, we generated Chr2-cl and Chr4-cl KO mice via ES cell injection into blastocysts, and after germline transmission we genotyped the offspring of heterozygous breeding pairs. While the offspring of Chr4-cl KO/WT parents were born close to Mendelian ratios in pure C57BL/6 and mixed C57BL/6 129Sv matings, one Chr4-cl KO/WT breeding pair gave birth to significantly fewer KO mice than expected (P value = 0.022) (Supplemental Fig. 5A). Likewise, two out of four Chr2-cl KO/WT breeding pairs on mixed C57BL/6 129Sv matings failed to give birth to a single KO/KO offspring (P value < 0.01) while the two other mating pairs produced KO/KO offspring at near Mendelian ratios (Supplemental Fig. 5A). Altogether, these data indicate that KRAB-ZFP clusters are not absolutely essential in mice, but that genetic and/or epigenetic factors may contribute to reduced viability.

We reasoned that retrotransposon activation could account for the reduced viability of Chr2-cl and Chr4-cl KO mice in some matings. However, since only rare matings produced non-viable KO embryos, we instead turned to the viable KO mice to assay for increased transposon activity. RNA-seq in blood, brain and liver revealed that, with a few exceptions, retrotransposons upregulated in KRAB-ZFP cluster KO ES cells are not expressed at higher levels in adult tissues, indicating that Chr2-cl and Chr4-cl KRAB-ZFPs are primarily required for TE repression during early development (Supplemental Fig. 5B). This is consistent with the high expression of these KRAB-ZFPs uniquely in ES cells (Supplemental Fig. 1A). To determine whether retrotransposition occurs at a higher frequency in Chr4-cl KO mice during development, we screened for novel ETn (ETn/ETnERV) and MuLV (MuLV/RLTR4_MM) insertions in viable Chr4-cl KO mice. For this purpose, we developed a capture-sequencing approach to enrich for ETn/RLTR4 DNA and flanking sequences from genomic DNA using probes that hybridize with the 5’ and 3’ ends of ETn and RLTR4 LTRs prior to deep sequencing. We screened genomic DNA samples from a total of 76 mice, including 54 mice from ancestry-controlled Chr4-cl KO matings in various strain backgrounds, the two ES cell lines the Chr4-cl KO mice were generated from, and eight mice from a Chr2-cl KO mating which served as a control (since ETn and MuLVs are not activated in Chr2-cl KO ESCs) (Supplemental Table 6). Using this approach, we were able to enrich reads mapping to ETn/MuLV LTRs about 2,000-fold compared to genome sequencing without capture. ETn/MuLV insertions were determined by counting uniquely mapped reads that were paired with reads mapping to ETN/MuLV elements (see materials and methods for details). Using this data, we first confirmed the polymorphic nature of both ETn and RLTR4 retrotransposons in laboratory mouse strains (Supplemental Fig. 6A-6B), highlighting the potential of these elements to retrotranspose. To identify novel insertions, we filtered out insertions that were supported by ETn/MuLV-paired reads in more than one animal. While none of the 54 ancestry-controlled mice showed a single novel MuLV insertion, we observed greatly varying numbers of up to 80 novel ETn insertions in our pedigree (Fig.4A).

To validate some of the novel ETn insertions in Chr4-cl KO mice, we designed specific PCR primers for five of the insertions and screened genomic DNA of the mice in which they were identified as well as their parents. For all tested insertions, we were able to amplify their flanking sequence and show that these insertions are absent in their parents (Supplemental Fig. 7A). To confirm the identity of these elements, we further amplified and sequenced three of the novel full-length ETn insertions (Genbank accession: MH449667-68). Based on the lower number of reads that supported novel ETn insertions relative to insertions inherited from the parents, we speculated that these elements retrotransposed in the developing embryo and not in the zygote or germ cells. Indeed, we detected different sets of insertions in various tissues from the same animal. Even between tail samples that were collected from the same animal at different ages, only a fraction of the new insertions were present in both samples, while technical replicates from the same genomic DNA samples showed a nearly complete overlap in insertions (Supplemental Fig. 7C). Besides novel somatic ETn insertions, we also observed three germ line retrotransposition events, as indicated by the presence of novel ETn insertions at identical locations in several siblings. One of these germ line insertions was also passed on to the next generation (Fig. 4A). Notably, these germ line insertions were supported by a significantly higher number of reads than the putative somatic insertions (Supplemental Fig. 7D).

Finally, we asked whether there were more ETn insertions in mice lacking the Chr4-cl relative to their wild type and heterozygous littermates in our pedigree. Interestingly, only one out of the eight Chr4-cl KO mice in a pure C56BL/6 strain background and none of the eight offspring from a Chr2-cl mating carried a single novel ETn insertion (Fig. 4A). When crossing into a 129Sv background for a single generation before intercrossing heterozygous mice (F1), we observed 4 out of 8 Chr4-cl KO/KO mice that contained at least one somatic ETn insertion, whereas none of 3 heterozygous mice contained any insertions. Upon crossing to the 129Sv background for a second generation (F2), we found that a similar fraction of wild type/heterozygous and KO/KO mice contained somatic ETn insertions, but that KO mice displayed a greater average number of insertions per animal (26 vs. 13, p= 0.057, Figure 4B). However, there was high variability in the number of insertions even amongst wild type mice. These data suggest that the Chr4-Cl KRAB-ZFPs may have a modest effect on ETn retrotransposition rates in some strains of mice, but other genetic and epigenetic effects also play an important role.

## Discussion

C2H2 zinc finger proteins, about half of which contain a KRAB repressor domain, represent the largest DNA-binding protein family in mammals. Nevertheless, most of these factors have not been investigated using loss-of-function studies. The most comprehensive characterization of human KRAB-ZFPs revealed a strong preference to bind TEs (*8*)(*11*) yet their function remains unknown. In humans, very few TEs are capable of retrotransposition yet many of them, often tens of million years old, are bound by KRAB-ZFPs. While this suggests that human KRAB-ZFPs mainly serve to control TE-borne enhancers and may have potentially transcription-independent functions, we were interested in the biological significance of KRAB-ZFPs in restricting potentially active TEs. The mouse is an ideal model for such studies since the mouse genome contains several active TE families, including IAP, ETn and L1 elements. We found that many of the young KRAB-ZFPs present in the genomic clusters of KRAB-ZFPs on chromosomes 2 and 4, which are highly expressed in a restricted pattern in ESCs, bound redundantly to these three active TE families. This provides strong evidence that many young KRAB-ZFPs are indeed expanding in response to TE activity. But do these young KRAB-ZFP genes limit the mobilization of TEs? Despite the large number of polymorphic ETn elements in mouse strains (*35*) and several reports of phenotype-causing novel ETn germ line insertions, no new ETn insertions were reported in a recent screen of 85 C57BL/6 mouse genomes (*36*), indicating that the overall rate of ETn germ line mobilization in inbred mice is rather low. We have demonstrated that Chr4-cl KRAB-ZFPs control ETn/ETnERV expression in ES cells, but this does not lead to widespread ETn mobility in viable C57BL/6 mice. In contrast, we found numerous somatic and several germ-line insertions in both WT and Chr4-cl KO mice in a C57BL/6 129Sv mixed genetic background, with generally more insertions in KO mice and in mice with more 129Sv DNA. Notably, there was a large variation in the number of new insertions in these mice, possibly caused by hyperactive polymorphic ETn insertions that varied from individual to individual, epigenetic variation at ETn insertions between individuals and/or the general stochastic nature of ETn mobilization. Furthermore, recent reports have suggested that KRAB-ZFP gene content is distinct in different strains of laboratory mice (*37, 38*), and reduced KRAB-ZFP gene content could contribute to increased activity in individual mice. Although we have yet to find obvious phenotypes in the mice carrying new insertions, novel ETn germ line insertions have been shown to cause phenotypes from short tails (*39-41*) to limb malformation (*3*) and severe morphogenetic defects including polypodia (*42*) depending upon their insertion site.

Despite a lack of widespread ETn activation in Chr4 cl KO mice, it still remains to be determined whether other TEs, like LINE1, IAP or other LTR retrotrotransposons are activated in any of the KRAB-ZFP cluster KO mice, which will require the development of additional capture-seq based assays. Notably, two of the heterozygous matings from Chr2 cl-KO mice failed to produce viable knockout offspring, which could indicate a TE-reactivation phenotype. It may also be necessary to generate compound homozygous mutants of distinct KRAB-ZFP clusters to eliminate redundancy before TEs become unleashed. The KRAB-ZFP cluster knockouts produced here will be useful reagents to test such hypotheses. In sum, our data supports that a major driver of KRAB-ZFP gene expansion in mice is recent retrotransposon insertions, and that redundancy within the KRAB-ZFP gene family and with other TE restriction pathways provides protection against widespread TE mobility, explaining the non-essential function of the majority of KRAB-ZFP genes.

## Materials and methods

### Cell lines and transgenic mice

Mouse ES cells and F9 EC cells were cultivated as described previously (*24*) unless stated otherwise. Chr4-cl KO ES cells originate from B6;129-Gt(ROSA)26Sortm1(cre/ERT)Nat/J mice (Jackson lab), all other KRAB-ZFP cluster KO ES cell lines originate from JM8A3.N1 C57BL/6N-A^tm1Brd^ ES cells (KOMP Repository). Chr2-cl KO and WT ES cells were initially grown in serum-containing media (*24*) but changed to 2i media (*43*) for several weeks before analysis. To generate Chr4-cl and Chr2-cl KO mice, the cluster deletions were repeated in B6 ES or R1 ES cells, respectively, and heterozygous clones were injected into B6 albino blastocysts. Chr2-cl KO mice were therefore kept on a mixed B6×129/Svx129/Sv-CP strain background while Chr4-cl KO mice were initially derived on a pure C57BL/6 background. For capture-seq screens, Chr4-cl KO/KO mice were crossed with 129X1/SvJ mice (Jackson lab) to produce the founder mice for Chr4-cl KO/KO and WT (B6/129 F1) offspring. Chr4-cl KO/+ (B6/129 F1) were also crossed with 129X1/SvJ mice to get Chr4-cl KO/+ (B6/129 F1) mice, which were intercrossed to give rise to the parents of Chr4-cl KO/KO and KO/+ (B6/129 F2) offspring.

### Generation of KRAB-ZFP expressing cell lines

KRAB-ZFP ORFs were PCR-amplified from cDNA or synthesized with codon-optimization (Supplemental Table 1), and stably expressed with 3XFLAG or 3XHA tags in F9 EC or ES cells using the *Sleeping beauty* transposon-based (*24*) or lentiviral expression vectors (Supplemental Table 1). Cells were selected with puromycin (1 µg/ml) and resistant clones were pooled and further expanded for ChIP-seq.

### CRISPR/Cas9 mediated deletion of KRAB-ZFP clusters and an MMETN insertion

All gRNAs were expressed from the pX330-U6-Chimeric_BB-CBh-hSpCas9 vector (Addgene #42230) and nucleofected into 10^6^ ES cells using Amaxa nucleofection in the following amounts: 5 µg of each pX330-gRNA plasmid, 1 µg pPGK-puro and 500 pmoles single-stranded repair oligos (Supplemental Table5). One day after nucleofection, cells were kept under puromycin selection (1 µg/ml) for 24 hours. Individual KO and WT clones were picked 7-8 days after nucleofection and expanded for PCR genotyping (Supplemental Table 5).

### ChIP-seq analysis

For ChIP-seq analysis of KRAB-ZFP expressing cells, 5-10 × 10^7^ cells were crosslinked and immunoprecipitated with anti-FLAG (Sigma, F1804) or anti-HA (Abcam, ab9110 or Covance, clone 16B12) antibody using one of two previously described protocols (*8, 44*) as indicated in Supplemental Table 1. H3K9me3 distribution in Chr4-cl, Chr10-cl, Chr13.1-cl and Chr13.2-cl KO ES cells was determined by native ChIP-seq and anti-H3K9me3 serum (Active Motif, #39161) as described previously (*45*). In Chr2-cl KO ES cells, H3K9me3 and KAP1 ChIP-seq was performed as previously described (*19*). In Chr4-cl KO and WT ES cells KAP1 binding was determined by endogenous tagging of KAP1 with C-terminal GFP (Supplemental Table 5), followed by FACS to enrich for GFP-positive cells and ChIP with anti-GFP (ThermoFisher, A-11122) using a previously described protocol (*44*). For ChIP-seq analysis of active histone marks, cross-linked chromatin from ES cells or testis (from two-week old mice) was immunoprecipitated with antibodies against H3K4me3 (ab8580), H3K4me1 (ab8895) and H3K27ac (ab4729) following the protocol developed by O’Geen *et al.* (*44*) or Khil *et al.* (*46*) respectively.

ChIP-seq libraries were constructed and sequenced as indicated in Supplemental Table 6. Reads were mapped to the mm9 genome using Bowtie (--best) or Bowtie2 as indicated in Supplemental Table 6 and peaks were called using MACS14 under high stringency settings (*P* < 1e-10, peak enrichment > 20) (*47*). Peaks were called both over the Input control and a FLAG or HA control ChIP (unless otherwise stated in Supplemental Table 6) and only peaks that were called in both settings were kept for further analysis. In cases when the stringency settings did not result in at least 50 peaks, the settings were changed to medium (*P* < 1e-10, peak enrichment > 10) or low (*P* < 1e-5, peak enrichment > 10) stringency (Supplemental Table 6). For further analysis, all peaks were scaled to 200 bp regions centered around the peak summits. The overlap of the scaled peaks to each repeat element (UCSC, mm9) were calculated by using the bedfisher function from BEDTools, with the setting -f 0.25. The right-tailed p-values between pair-wise comparison of each ChIP-Seq peak and repeat element were extracted, and then adjusted using the Benjamini-Hochberg approach implemented in the R function p.adjust(). Binding motifs were determined using only nonrepetitive (< 10% repeat content) peaks with MEME (*48*). MEME motifs were compared with *in silico* predicted motifs (*11*) using Tomtom (*48*) and considered as significantly overlapping with a False Discovery Rate (FDR) below 0.1. To find MEME and predicted motifs in repetitive peaks, we used FIMO (*48*). Differential H3K9me3 and KAP1 distribution in WT and Chr2-cl or Chr4-cl KO ES cells at TEs was determined by counting ChIP-seq reads overlapping annotated insertions of each TE group using BEDTools (MultiCovBed). Additionally, ChIP-seq reads were counted at the TE fraction that was bound by Chr2-cl or Chr4-cl KRAB-ZFPs (overlapping with 200 bp peaks). Count tables were concatenated and analyzed using DESeq2 (*49*). The previously published ChIP-seq datasets for KAP1 (*50*) and H3K9me3 (*51*) were re-mapped using Bowtie (--best).

### Luciferase reporter assays

For KRAB-ZFP repression assays, double-stranded DNA oligos containing KRAB-ZFP target sequences (Supplemental Table 5) were cloned upstream of the SV40 promoter of the pGL3-Promoter vector (Promega) between the restriction sites for NheI and XhoI. 33 ng of reporter vectors were co-transfected (Lipofectamine 2000, Thermofisher) with 33 ng pRL-SV40 (Promega) for normalization and 33 ng of transient KRAB-ZFP expression vectors (in pcDNA3.1) or empty pcDNA3.1 into 293T cells seeded one day earlier in 96-well plates. Cells were lysed 48 hours after transfection and luciferase/Renilla luciferase activity was measured using the Dual-Luciferase Reporter Assay System (Promega). To measure the transcriptional activity of the MMETN element upstream of the *Cd59a* gene, fragments of varying sizes (Supplemental Table 5) were cloned into the promoter-less pGL3-basic vector (Promega) using NheI and NcoI sites. 70 ng of reporter vectors were cotransfected with 30 ng pRL-SV40 into feeder-depleted Chr4-cl WT and KO ES cells, seeded into a gelatinized 96 well plate 2 hours before transfection. Luciferase activity was measured 48 hours after transfection as described above.

### RNA-seq analysis

Whole RNA was purified using RNeasy columns (Qiagen) with on column DNase treatment or the High Pure RNA Isolation Kit (Roche) (Supplemental Table 6). Tissues were first lysed in TRIzol reagent (ThermoFisher) and RNA was purified after the Isopropanol precipitation step using RNeasy columns (Qiagen) with on column DNase treatment. Libraries were generated using the SureSelect Strand-Specific RNA Library Prep kit (Agilent) or Illumina’s TruSeq RNA Library Prep Kit (with polyA selection) and sequenced as 50 or 100 bp paired-end reads on an Illumina HiSeq2500 or HiSeq3000 machine (Supplemental Table 6). RNA-seq reads were mapped to the mouse genome (mm9) using Tophat (--I 200000 --g 1) unless otherwise stated. For differential Transposon expression, mapped reads that overlap with TE elements annotated in the Repeatmasker track (http://www.repeatmasker.org) were counted using BEDTools MultiCovBed (-split). Reads mapping to multiple fragments that belong to the same TE insertion (as indicated by the repeat ID) were summed up. Only transposons with a total of at least 20 (for two biological replicates) or 30 (for three biological replicates) mapped reads across WT and KO samples were considered for differential expression analysis. Transposons within the deleted KRAB-ZFP cluster were excluded from the analysis. Read count tables were used for differential expression analysis with DESeq2. For differential gene expression analysis, reads overlapping with gene exons were counted using HTSeq-count and analyzed using DESeq2. To test if KRAB-ZFP peaks are significantly enriched near up-or down-regulated genes, a binomial test was performed. Briefly, the proportion of the peaks that are located within a certain distance up-or downstream to the TSS of genes was determined using the windowBed function of BED tools. The probability *p* in the binomial distribution was estimated as the fraction of all genes overlapped with KRAB-ZFP peaks. Then, given *n* which is the number of specific groups of genes, and *x* which is the number of this group of genes overlapped with peaks, the R function binom.test() was used to estimate the p-value based on right-tailed Binomial test. Finally, the adjusted p-values were determined separately for LTR and LINE retrotransposon groups using the Benjamini-Hochberg approach implemented in the R function p.adjust().

### Reduced Representation Bisulfite sequencing (RRBS-seq)

For RRBS-seq analysis, Chr4-cl WT and KO ES cells were grown in either standard ES cell media containing FCS or for one week in 2i media containing vitamin C as described previously (*32*). Genomic DNA was purified from WT and Chr4-cl KO ES cells using the Quick-gDNA purification kit (Zymo Research) and bisulfite-converted with the NEXTflex™ Bisulfite-Seq Kit (Bio Scientific) using Msp1 digestion to fragment DNA. Libraries were sequenced as 50bp paired-end reads on an Illumina HiSeq. The reads were processed using Trim Galore (--illumina --paired –rrbs) to trim poor quality bases and adaptors. Additionally, the first 5 nt of R2 and the last 3 nt of R1 and R2 were trimmed. Reads were then mapped to the reference genome (mm9) using Bismark (*52*) to extract methylation calling results. The averaged CpG methylation for each repeat family was calculated using a custom script. For comparison of CpG methylation between WT and Chr4-cl KO ES cells (in serum or 2i + Vitamin C conditions) only CpG sites with at least 10-fold coverage in each sample were considered for analysis.

### Retrotransposition assay

The retrotransposition vectors pCMV-MusD2, pCMV-MusD2-neoTNF and pCMV-ETnI1-neoTNF (*30*) were a kind gift from Dixie Mager. To partially delete the Gm13051 binding site within pCMV-MusD2-neoTNF, the vector was cut with KpnI and re-ligated using a repair oligo, leaving a 24 bp deletion within the Gm13051 binding site. The Rex2 binding site in pCMV-ETnI1-neoTNF was deleted by cutting the vector with EcoR1 and XbaI followed by re-ligation using two overlapping PCR products, leaving a 45 bp deletion while maintaining the rest of the vector unchanged (see Supplemental Table 5 for primer sequences). For MusD retrotransposition assays, 5×10^4^ HeLa cells were transfected in a 24-well dish with 100 ng pCMV-MusD2-neoTNF or pCMV-MusD2-neoTNF (ΔGm13051-m) using Lipofectamine 2000. For ETn retrotransposition assays, 50 ng of pCMV-ETnI1-neoTNF or pCMV-ETnI1-neoTNF (ΔRex2) vectors were cotransfected with 50 ng pCMV-MusD2 to provide gag and pol proteins in trans. G418 (0.6 mg/ml) was added five days after transfection and cells were grown under selection until colonies were readily visible by eye. G418-resistant colonies were stained with Amido Black (Sigma).

### Capture-seq screen

To identify novel retrotransposon insertions, genomic DNA from various tissues (Supplemental Table 6) was purified and used for library construction with target enrichment using the SureSelect^QXT^ Target Enrichment kit (Agilent). Custom RNA capture probes were designed to hybridize with the 120 bp 5’ ends of the 5’LTRs and the 120 bp 3’ ends of the 3’LTR of about 600 intact (internal region flanked by two LTRs) MMETn/RLTRETN retrotransposons or of 140 RLTR4_MM/RLTR4 retrotransposons that were upregulated in Chr4-cl KO ES cells. Enriched libraries were sequenced on an Illumina HiSeq as paired-end 50bp reads. R1 and R2 reads were mapped to the mm9 genome separately, using settings that only allows non-duplicated, uniquely mappable reads (Bowtie -m 1 --best --strata; samtools rmdup -s) and under settings that allows multimapping and duplicated reads (Bowtie --best). Of the latter, only reads that overlap (min. 50% of read) with RLTRETN, MMETn-int, ETnERV-int, ETnERV2-int or ETnERV3-int repeats (ETn) or RLTR4, RLTR4_MM-int or MuLV-int repeats (RLTR4) were kept. Only uniquely mappable reads whose paired reads were overlapping with the repeats mentioned above were used for further analysis. All ETn- and RLTR4-paired reads were then clustered (as bed files) using BEDTools (bedtools merge -i -n -d 1000) to receive a list of all potential annotated and non-annotated new ETn or RLTR4 insertion sites and all overlapping ETn- or RLTR4-paired reads were counted for each sample at each locus. Finally, all regions that were located within 1 kb of an annotated RLTRETN, MMETn-int, ETnERV-int, ETnERV2-int or ETnERV3-int repeat as well as regions overlapping with previously identified polymorphic ETn elements (*35*) were removed. Genomic loci with at least 10 reads per million unique ETn- or RLTR4-paired reads were considered as insertion sites. To qualify for a de-novo insertion, we allowed no called insertions in any of the other screened mice at the locus and not a single read at the locus in the ancestors of the mouse. Insertions at the same locus in at least two siblings from the same offspring were considered as germ-line insertions, if the insertion was absent in the parents and mice who were not direct descendants from these siblings. Full-length sequencing of new ETn insertions was done by Sanger sequencing of short PCR products in combination with Illumina sequencing of a large PCR product (Supplemental Table 5), followed by de-novo assembly using the Unicycler software.

## Data and materials availability

All NGS data has been deposited in GEO (GSE115291). Sequences of full-length *de novo* ETn insertions have been deposited in the GenBank database (MH449667-MH449669).

## Acknowledgments

We thank Alex Grinberg, Jeanne Yimdjo and Victoria Carter for generating and maintaining transgenic mice. We also thank members of the Macfarlan and Trono labs for useful discussion, Steven Coon, James Iben, Tianwei Li and Anna Malawska for NGS and computational support. This work was supported by NIH grant 1ZIAHD008933 and the NIH DDIR Innovation Award program (T.S.M.), and by subsidies from the Swiss National Science Foundation (310030_152879 and 310030B_173337) and the European Research Council (KRABnKAP, No. 268721; Transpos-X, No. 694658) (DT).

## Author contributions

G.W., A.D., T.S.M. and D.T. conceived the study, G.W., A.D., M.B., M.T., D.H. and S.R. performed the experiments, G.W., A.D., M.S. and A.M. analyzed the data.

---

**Supplemental Figure 1:**
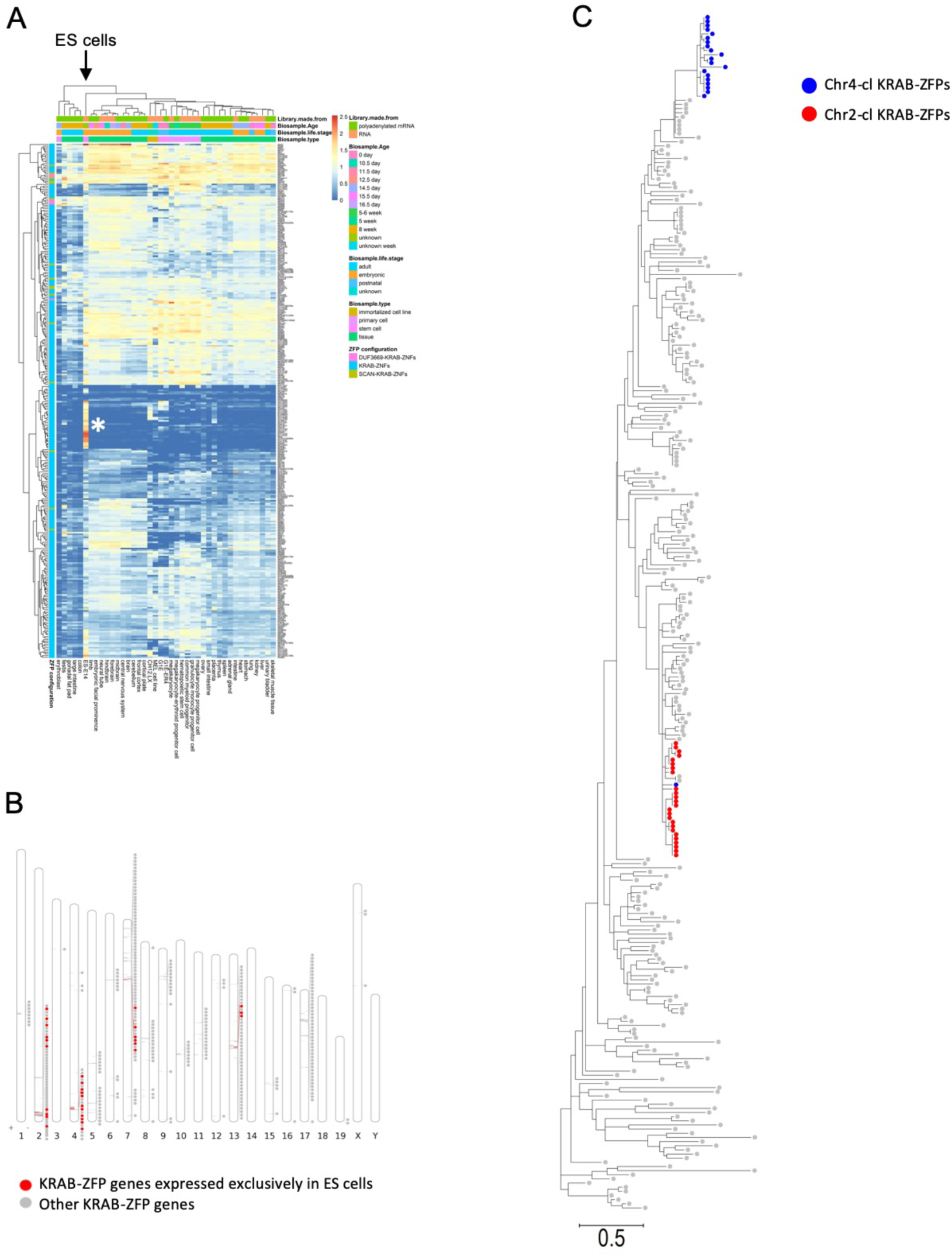
ES cell-specific expression of KRAB-ZFP gene clusters. (A) Heatmap showing expression patterns of mouse KRAB-ZFPs in 40 mouse tissues and cell lines (ENCODE). Heatmap colors indicate gene expression levels in transcripts per million (TPM). The asterisk indicates a group of 30 KRAB-ZFPs that are exclusively expressed in ES cells. (B) Physical location of the genes encoding for the 30 KRAB-ZFPs that are exclusively expressed in ES cells. (C) Phylogenetic (Maximum likelihood) tree of the KRAB domains of mouse KRAB-ZFPs. KRAB-ZFPs encoded on the gene clusters on chromosome 2 and 4 are highlighted.

**Supplemental Figure 2:**
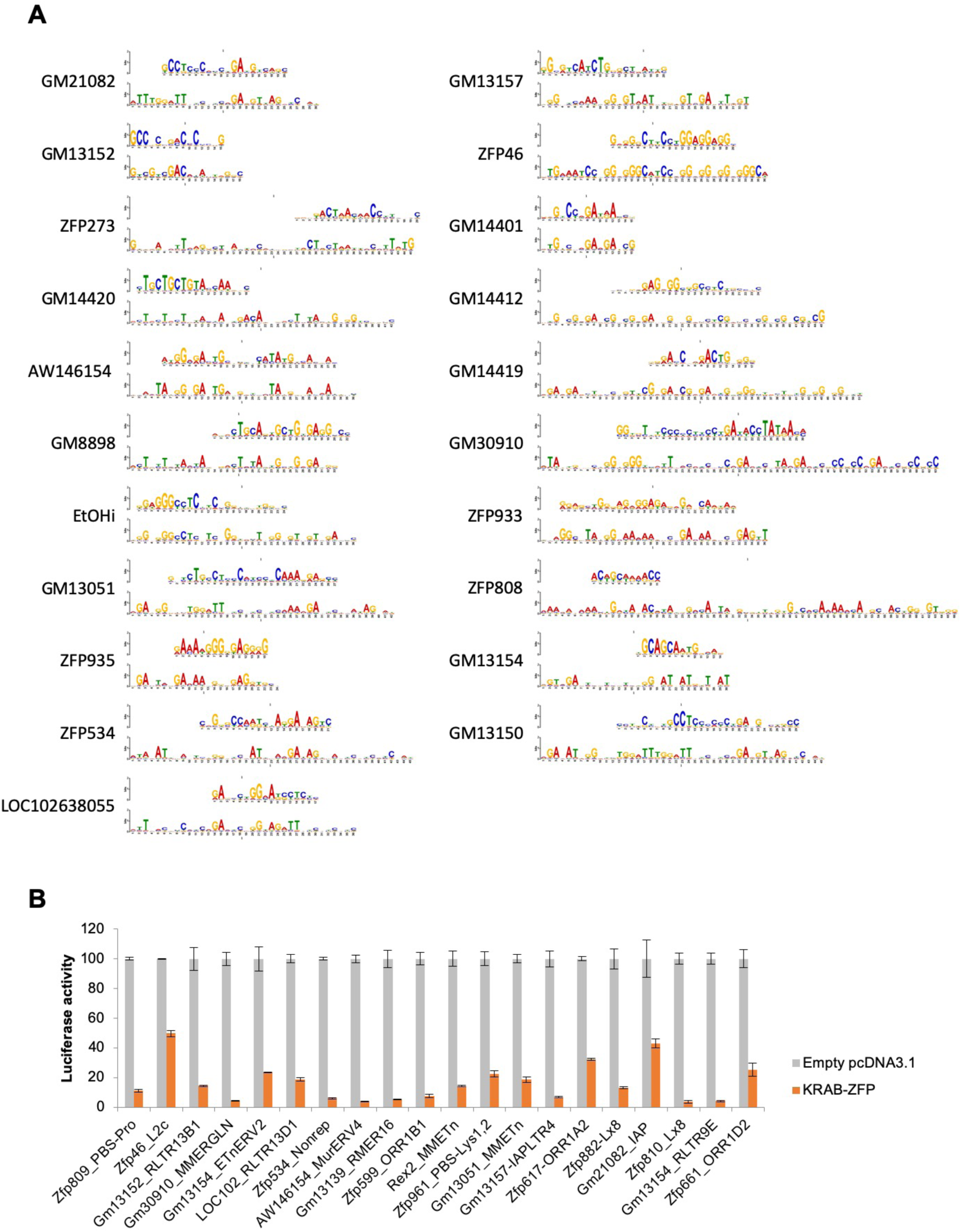
KRAB-ZFP binding motifs and their repression activity. (A) Comparison of computationally predicted (bottom) and experimentally determined (top) KRAB-ZFP binding motifs. Only significant pairs are shown (FDR < 0.1). (B) Luciferase reporter assays to confirm KRAB-ZFP repression of the identified target sites. Bars show the luciferase activity (normalized to Renilla luciferase) of reporter plasmids containing the indicated target sites cloned upstream of the SV40 promoter. Reporter plasmids were co-transfected into 293T cells with a Renilla luciferase plasmid for normalization and plasmids expressing the targeting KRAB-ZFP. Normalized luciferase activity is shown relative to luciferase activity of the reporter plasmid co-transfected with an empty pCDNA3.1 vector.

**Supplemental Figure 3:**
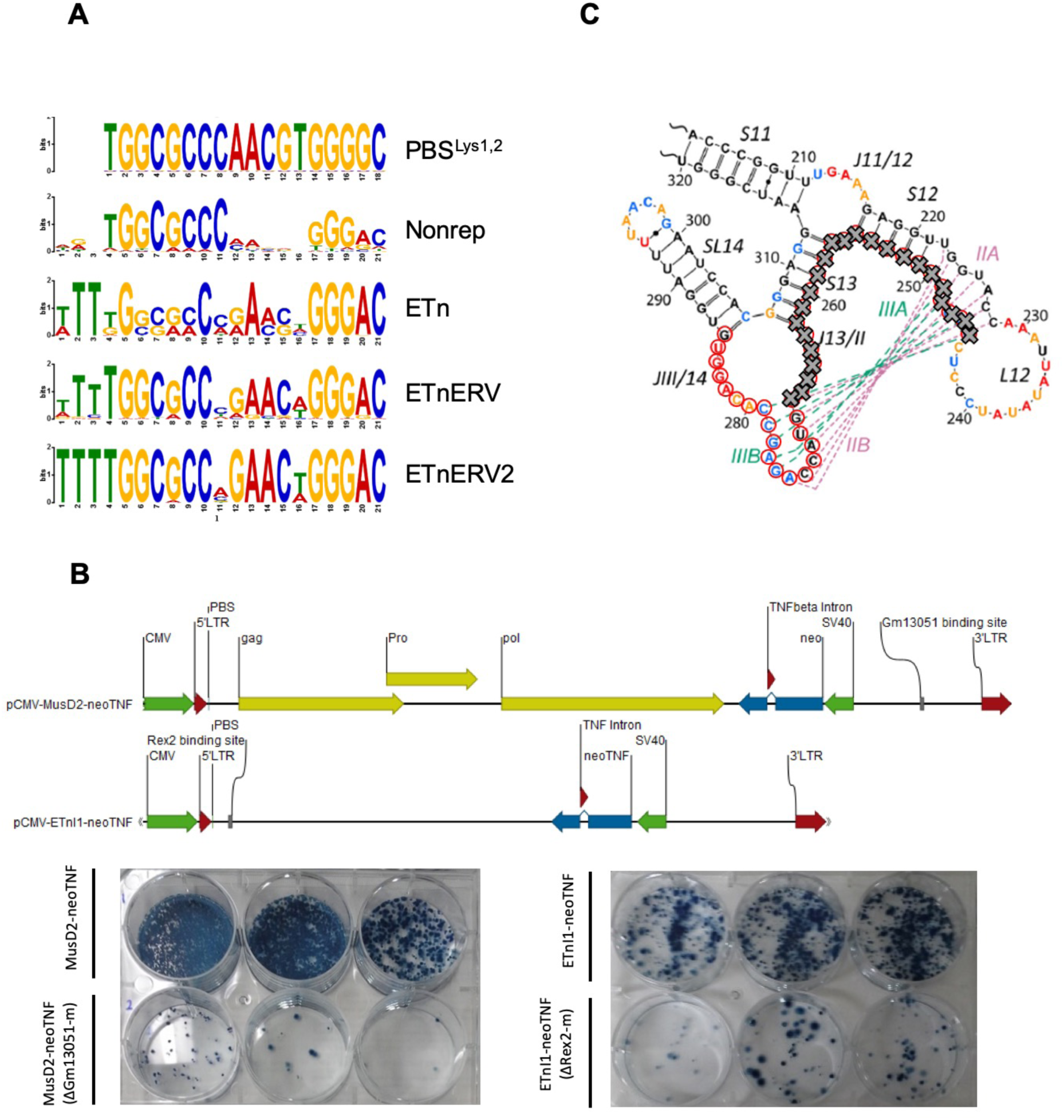
KRAB-ZFP binding to ETn retrotransposons. (A) Comparison of the PBS^Lys1,2^ sequence with Zfp961 binding motifs in nonrepetitive peaks (Nonrep) and peaks at ETn elements. (B) Retrotransposition assays of original (ETnI1-neoTNF and MusD2-neoTNF (*30*)) and modified reporter vectors where the Rex2 or Gm13051 binding motifs where removed. Schematic of reporter vectors are siplayed at the top. HeLa cells were transfected as described in the Materials and Methods section and neo-resistant colonies, indicating retrotransposition events, were selected and stained. (C) Stem-loop structure of the ETn RNA export signal, the Gm13051 motif on the corresponding DNA is marked with red circles, the part of the motif that was deleted is indicated with grey crosses (adapted from ref (*31*)).

**Supplemental Figure 4:**
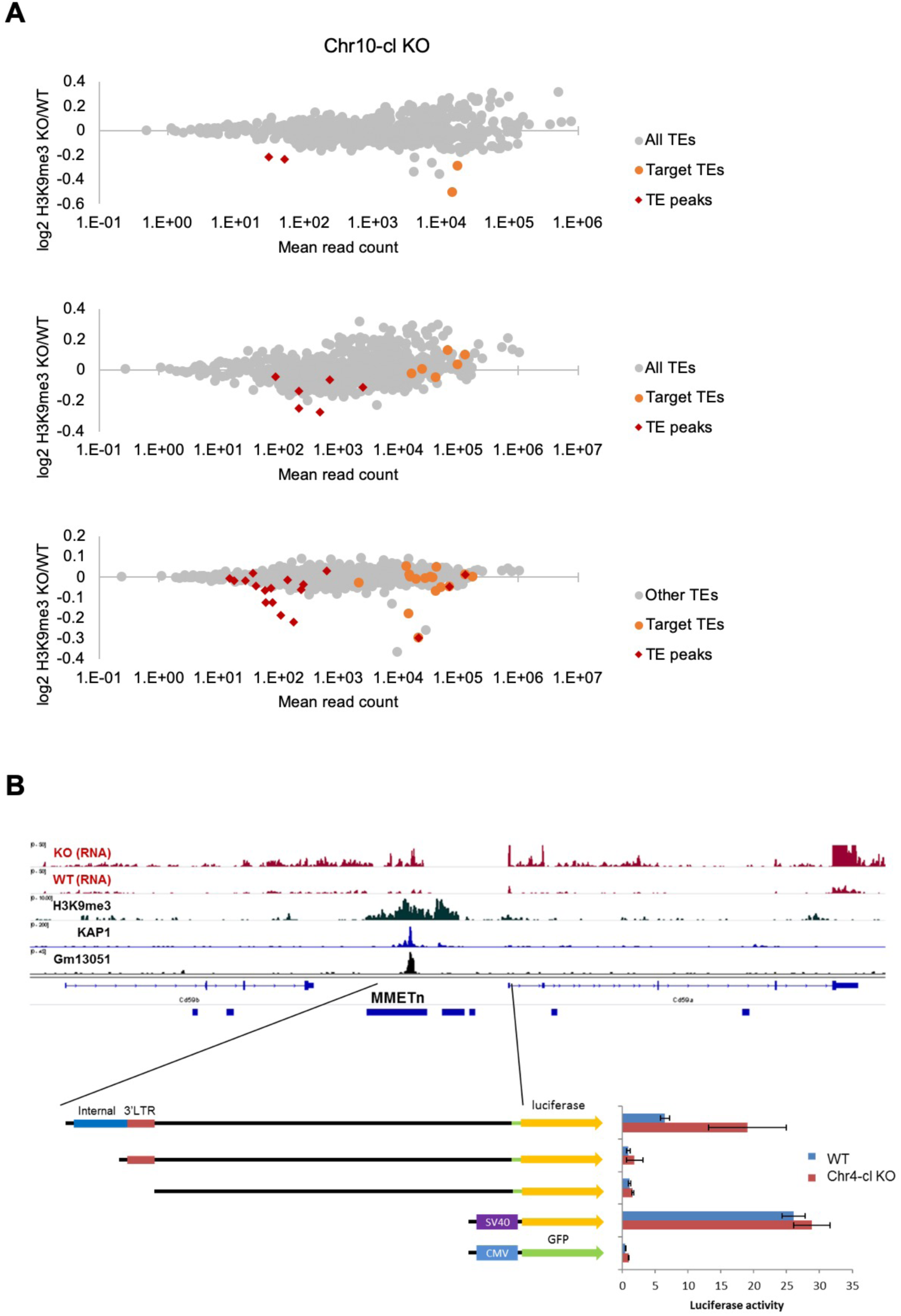
(A) Differential analysis of summative (all individual insertions combined) H3K9me3 enrichment at TE groups in Chr10-cl, Chr13.1 and Chr13.2-cl KO ES cells. TE groups targeted by one or several KRAB-ZFPs encoded within the deleted clusters are highlighted in orange (differential enrichment over the entire TE sequences) and red (differential enrichment at TE regions that overlap with KRAB-ZFP ChIP-seq peaks). (B) Top: Schematic view of the *Cd59a*/*Cd59b* locus with a 5’truncated ETn insertion. ChIP-seq (Input subtracted from ChIP) data for overexpressed epitope-tagged Gm13051 (a Chr4-cl KRAB-ZFP) in F9 EC cells, and re-mapped KAP1 (GEO accession: GSM1406445) and H3K9me3 (GEO accession: GSM1327148) in WT ES cells are shown together with RNA-seq data from Chr4-cl WT and KO ES cells (mapped using Bowtie (-a -m 1 --strata -v 2) to exclude reads that cannot be uniquely mapped). Bottom: Transcriptional activity of a 5 kb fragment with or without fragments of the ETn insertion was tested by luciferase reporter assay in Chr4-cl WT and KO ES cells.

**Supplemental Figure 5:**
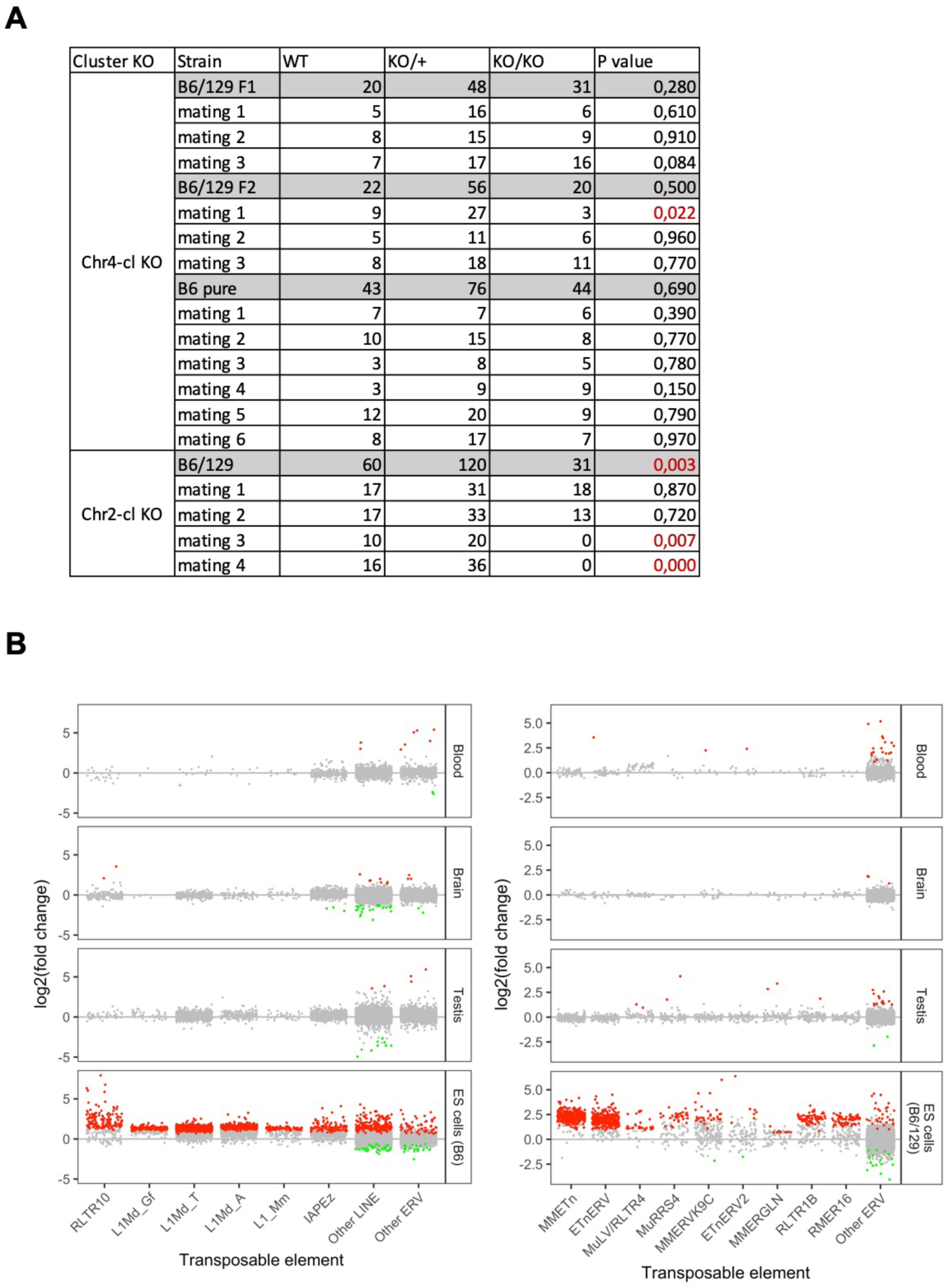
(A) Birth statistics of Chr4- and Chr2-cl mice derived from KO/WT x KO/WT matings in different strain backgrounds. (B) RNA-seq analysis of TE expression in Chr2- and Chr4-cl KO tissues. TE groups with the highest reactivation phenotype in ES cells are shown separately. Significantly up- and downregulated elements (*adjusted p value* < 0.05) are highlighted in red and green, respectively. Experiments were performed in at least two biological replicates.

**Supplemental Figure 6:**
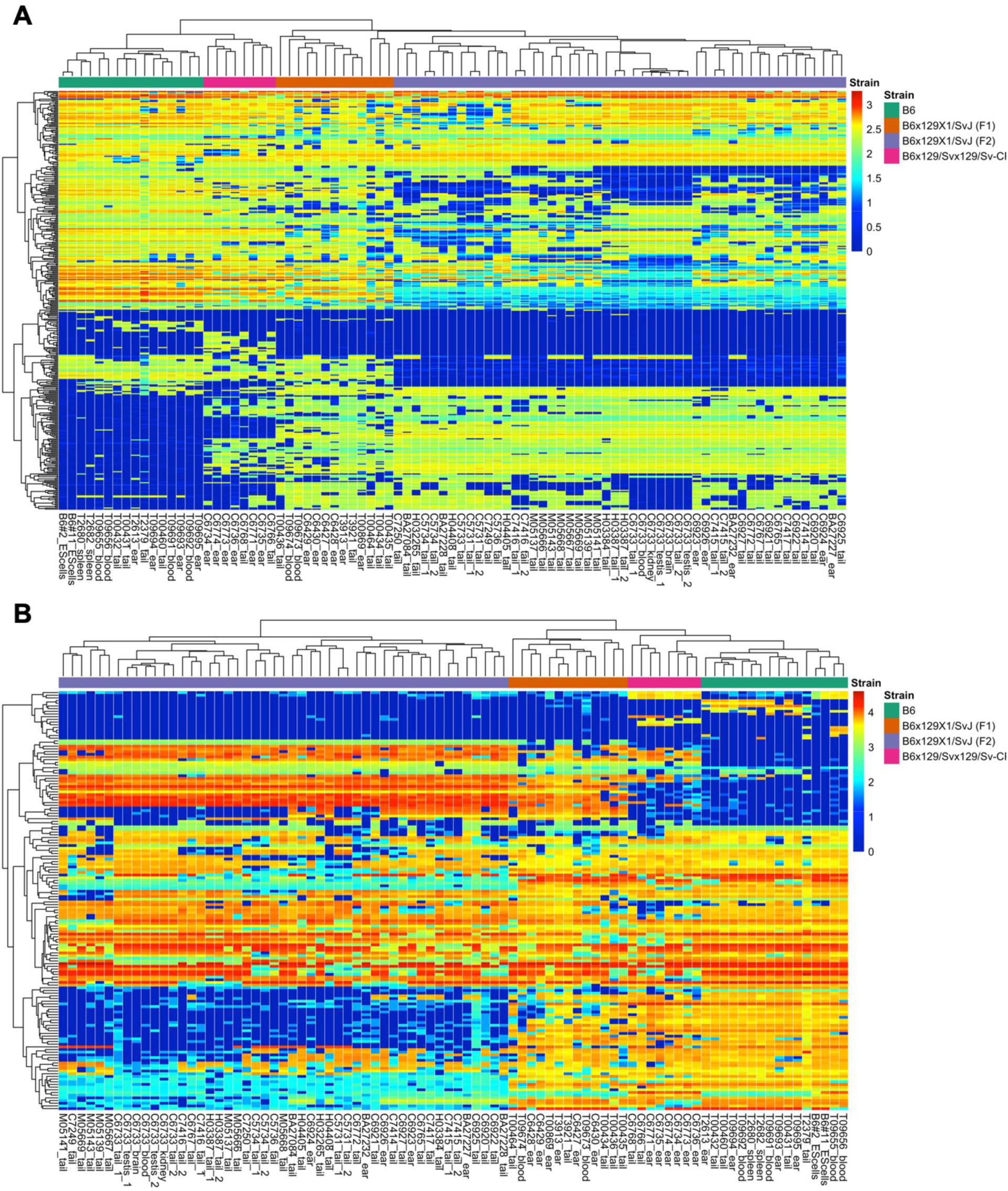
Identification of polymorphic ETn and MulV retrotransposon insertions in Chr4-cl KO and WT mice. Heatmaps show normalized Capture-Seq read counts in RPM (Read Per Million) for identified polymorphic ETn (A) and MulV (B) loci in different mouse strains. Only loci with strong support for germ-line ETn or MuLV insertions (at least 100 or 3000 ETn or MuLV RPM, respectively) in at least two animals are shown. Non-polymorphic insertion loci with high read counts in all screened mice were excluded for better visibility. The sample information (sample name and cell_type/tissue) is annotated at the bottom, with the strain information indicated by color at the top. The color gradient indicates log10(RPM+1).

**Supplemental Figure 7:**
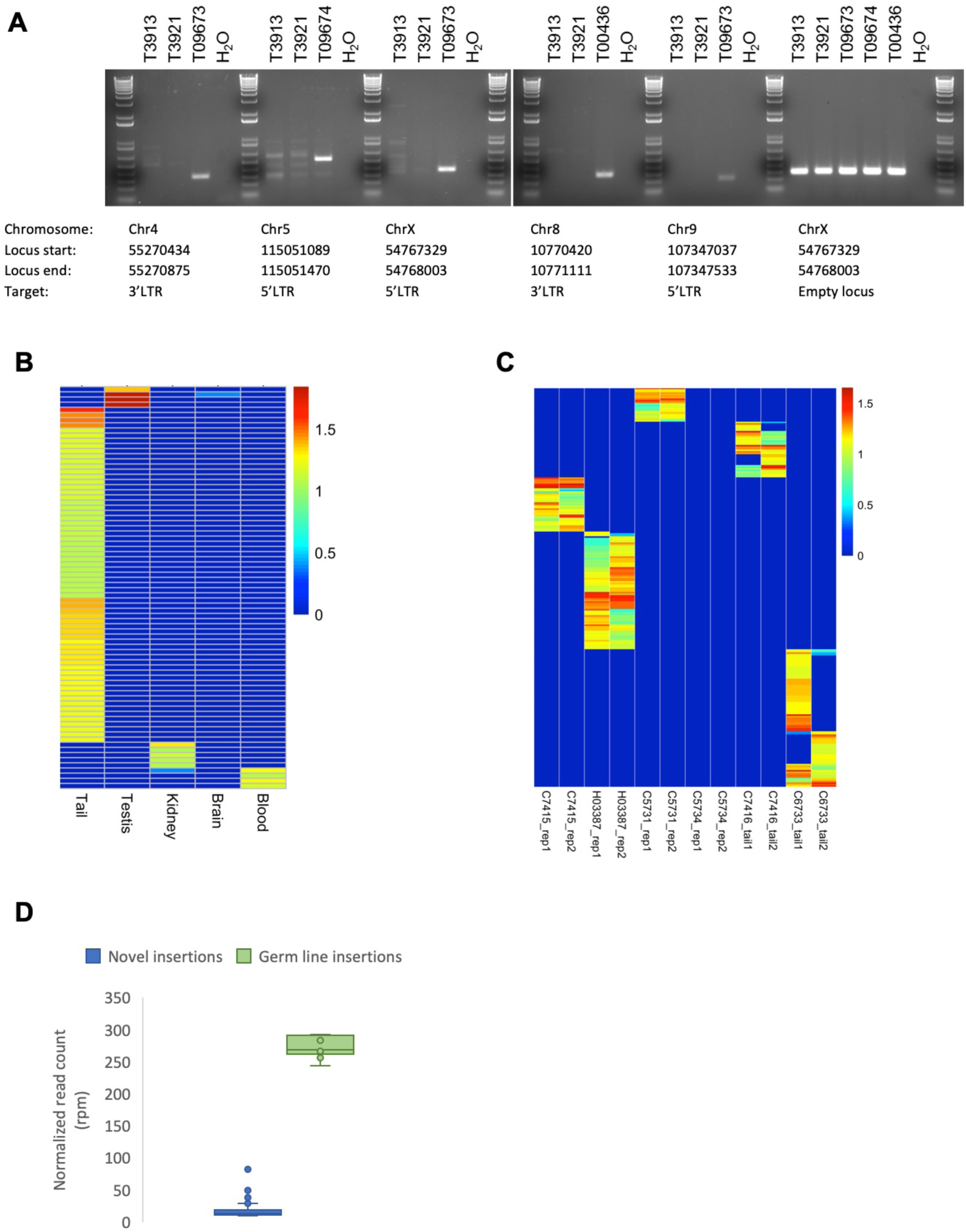
Confirmation of novel ETn insertions identified by capture-seq. (A) PCR confirmation of novel ETn insertions in genomic DNA of three littermates (IDs: T09673, T09674 and T00436) and their parents (T3913 and T3921). Primer sequences are shown in Table S5. (B) Heatmap shows Capture-Seq read counts (RPM) of a Chr4-cl KO mouse (ID: C6733) as determined in different tissues. Each row represents a novel ETn loci that was identified in at least one tissue. The color gradient indicates log10(RPM+1). (C) Heatmap shows the Capture-Seq RPM in technical replicates using the same Chr4-cl KO DNA sample (rep1/rep2) or replicates with DNA samples prepared from different sections of the tail from the same mouse at different ages (tail1/tail2). Each row represents a novel ETn loci that was identified in at least one of the displayed samples. The color gradient indicates log10(RPM+1). (D) ETn Capture-Seq read counts (RPM) at putative novel somatic (loci identified exclusively in one single animal) and germ-line (loci identified in several littermates) insertions.

